# Association between SARS-CoV-2 neutralizing antibodies and commercial serological assays

**DOI:** 10.1101/2020.07.01.182220

**Authors:** Mei San Tang, James Brett Case, Caroline E. Franks, Rita E. Chen, Neil W. Anderson, Jeffrey P. Henderson, Michael S. Diamond, Ann M. Gronowski, Christopher W. Farnsworth

**Affiliations:** Department of Pathology & Immunology. Washington University School of Medicine. St. Louis, MO; Department of Medicine. Washington University School of Medicine. St. Louis, MO; Department of Molecular Microbiology. Washington University School of Medicine. St. Louis, MO

**Author notes:** These authors contributed equally to this work (Mei San Tang, James Brett Case). Address for Correspondence Christopher W. Farnsworth, Department of Pathology & Immunology, Washington University in St. Louis, 660 S. Euclid Ave, Campus Box 8118, St. Louis, MO 63110. Drs’ Farnsworth and Gronowski are co-corresponding authors.

**Keywords:** Serology, Neutralizing antibodies, COVID-19, SARS-CoV-2

## Abstract

**Introduction:** Commercially available SARS-CoV-2 serological assays based on different viral antigens have been approved for the qualitative determination of anti-SARS-CoV-2 antibodies. However, there is limited published data associating the results from commercial assays with neutralizing antibodies.

**Methods:** 67 specimens from 48 patients with PCR-confirmed COVID-19 and a positive result by the Roche Elecsys SARS-CoV-2, Abbott SARS-CoV-2 IgG, or EUROIMMUN SARS-CoV-2 IgG assays and 5 control specimens were analyzed for the presence of neutralizing antibodies to SARS-CoV-2. Correlation, concordance, positive percent agreement (PPA), and negative percent agreement (NPA) were calculated at several cutoffs. Results were compared in patients categorized by clinical outcomes.

**Results:** The correlation between SARS-CoV-2 neutralizing titer (EC_50_) and the Roche, Abbott, and EUROIMMUN assays was 0.29, 0.47, and 0.46 respectively. At an EC_50_ of 1:32, the concordance kappa with Roche was 0.49 (95% CI; 0.23-0.75), with Abbott was 0.52 (0.28-0.77), and with EUROIMMUN was 0.61 (0.4-0.82). At the same neutralizing titer, the PPA and NPA for the Roche was 100% (94-100) & 56% (30-80); Abbott was 96% (88-99) & 69% (44-86); and EUROIMMUN was 91% (80-96) & 81% (57-93) for distinguishing neutralizing antibodies. Patients who died, were intubated, or had a cardiac injury from COVID-19 infection had significantly higher neutralizing titers relative to those with mild symptoms.

**Conclusion:** COVID-19 patients generate an antibody response to multiple viral proteins such that the calibrator ratios on the Roche, Abbott, and EUROIMMUN assays are all associated with SARS-CoV-2 neutralization. Nevertheless, commercial serological assays have poor NPA for SARS-CoV-2 neutralization, making them imperfect proxies for neutralization.

## INTRODUCTION

Host cell infections by the recently-emerged severe acute respiratory syndrome coronavirus 2 (SARS-CoV-2) begin when the viral spike (S) protein engages the host angiotensin-converting enzyme 2 (ACE2) receptor *(1)*. The humoral immune response can block infection through neutralizing antibodies, which bind the virus in a manner that prevents host cell infection *(2)*. For SARS-CoV-2, this may be achieved by interfering with the spike -ACE2 receptor interaction, or by disrupting the fusion mechanisms the virus uses to enter host cell cytoplasm *(2)*.

In the absence of a vaccine, there is considerable interest in identifying high-affinity neutralizing antibodies to SARS-CoV-2 to assess immune status and to evaluate vaccine responses. We previously demonstrated that passive transfer of monoclonal antibodies against SARS-CoV-2 S protein reduced viral titers and pathology in the lungs in a mouse model of SARS-CoV-2 *(3)*. Monoclonal antibodies engineered from neutralizing antibodies, initially identified from convalescent COVID-19 patients, have been advanced as potential antiviral therapeutics *(4-6)*, and early results from convalescent plasma use in patients indicate a protective effect of antibodies against SARS-CoV-2 *(7-10)*. While early results are promising, the antibody titer conferring protection remains unclear and the role of neutralizing antibodies in protection has not been fully elucidated *(11)*.

Despite widespread interest in neutralizing antibodies, methods for their detection and quantification are relatively low-throughput and limited to Biosafety Level 3-equipped research laboratories. While high-throughput methods have emerged, most rely on recombinant Vesicular Stomatitis Viruses (VSV) engineered to express a portion of the SARS-CoV-2 viral spike protein, and their subsequent entry into cell lines *(12-14)*. Commercially available serological assays are high-throughput, relatively inexpensive, and use readily available instrumentation. The use of automated serological SARS-CoV-2 assays as a surrogate for neutralizing titers is therefore an attractive option. To date, limited data are available correlating commercially available assays with the presence of neutralizing antibodies.

We previously compared the clinical performance of three commercial serological assays *(15, 16)*. Here, we further assess the ability of these assays to predict the presence of neutralizing antibodies.

## MATERIALS AND METHODS

### Specimens

This study was approved by the Institutional Review Board of Washington University in St. Louis. Residual plasma from physician-ordered complete blood count were utilized. Specimens were obtained from patients with PCR-confirmed COVID-19 and at least one previously positive SARS-CoV-2 serological result. A subset of pre-pandemic samples obtained in 2015 and stored at -80 °C were used as negative controls.

### Clinical information

Duration from symptom onset was obtained from two independent assessors by review of the electronic medical record (EMR) and inferred from physician encounter notes. Symptoms included cough, fever, shortness of breath, loss of taste or smell, sore throat, and headache *(17)*. The EMR also was used to collect data on outcomes for each patient. Mortality and intubation were determined by physician encounter notes, acute kidney injury (AKI) was defined using RIFLE criteria of 2-fold increase in serum creatinine and urine output less than 5 mL/kg/hr, cardiac injury was defined as a troponin I concentration > 0.03 ng/mL (Abbott Diagnostics).

### INSTRUMENTATION

Specimens were analyzed on three commercially available immunoassays and reported previously *(15, 16)*. The Roche Elecsys Anti-SARS-CoV-2 assay was performed on an a Cobas e 601. The Roche assay detects total antibodies (IgG, IgA, IgM) against an epitope of the viral nucleocapsid protein. The Abbott SARS-CoV-2 IgG assay was performed on an i2000 Abbott Architect (Abbott Diagnostics) and detects IgG antibodies against the viral nucleocapsid protein. The EUROIMMUN (EI) SARS-CoV-2 IgG assay was performed on a QUANTA-Lyser 240 (Inova Diagnostics) assay and detects anti-SARS-CoV-2 IgG directed against the S1 domain of viral spike protein. All three assays use an assay-specific calibrator to report the ratio of the signal from the specimen to the signal of the calibrator. The results are interpreted as positive or negative relative to a threshold value. For the Roche assay, a positive is a cutoff index (COI) ≥1; for the Abbott assay, a signal to cut-off (S/CO) ≥1.4 is positive and <1.4 is negative; for the EI assay, a ratio ≥ 1.2 is positive 0.80-1.19 is indeterminate, and < 0.8 is negative. The cutoff of 1.2 was used as a positive result for the EI. All three assays specify a positive result as the signal of the sample/the signal of a calibrator, therefore all results are reported here as a ratio.

### FOCUS REDUCTION NEUTRALIZATION ASSAYS

Neutralization assays were performed as previously described *(18)*. Briefly, SARS-CoV-2 strain 2019 n-CoV/USA_WA1/2020 was obtained from the Centers of Disease Control and passaged in Vero E6 cells with DMEM (Corning) supplemented with glucose, L-glutamine, sodium pyruvate, and 10% FBS. Indicated dilutions of plasma were incubated with 10^2^ focus forming units (FFU) of SARS-CoV-2 for 1h at 37°C before addition of the antibody virus complex to Vero E6 monolayers at 37°C for 1h. Cells were overlaid with a 1% w/v methylcellulose in MEM supplemented with 2% FBS and harvested 30h later. Methylcellulose overlays were removed and fixed with 4% paraformaldehyde in PBS at room temperature. Plates were then washed and incubated with 1 µg/mL anti-S antibody (CR3022) *(19)* and HRP-conjugated goat anti-Human IgG. Cells infected by SARS-CoV-2 were visualized using TrueBlue peroxidase substrate (KPL) and cell foci were quantified using an ImmunoSpot microanalyzer (Cellular Technologies). For each specimen, a minimum of 8 dilutions of human plasma were performed in duplicate and a standard curve generated. The 1/Log_10_ plasma dilution (EC_50_) is the dilution at which 50% of the cells were infected with virus and formed foci (**Supplemental Figure 1**).

### STATISTICS

Correlation between clinical assays and neutralizing titers were calculated using linear regression. Concordance between the assays was calculated using Cohen’s Kappa. Area under the curve (AUC) for receiver operator characteristic (ROC) curves were calculated using the Wilson/Brown method. Kappa, positive percent agreement (PPA), and negative percent agreement (NPA) analysis were performed using multiple cutoffs for neutralizing titers owing to a lack of consensus regarding the relevant protective titer. Differences between antibody and neutralizing titers categorized by outcomes were calculated using unpaired T-tests. For outcome comparisons, all specimens were >d10 post-symptom onset. All statistical analyses were performed with GraphPad Prism 8 (GraphPad).

## RESULTS

40/42 specimens from PCR-confirmed COVID-19 patients with positive antibody results from commercial SARS-CoV-2 assays had neutralizing titers >1:20 by d14 post-symptom onset (**Figure 1A**). The mean neutralizing titer by d21 was 1:250 (95% CI; 1:149-1:436). In contrast, pre-pandemic control samples were not neutralizing at a titer of 1:20. Neutralizing titers increased subsequently with days post-symptom onset (**Supplemental Figure 2**). A subset of patients with serial measurements demonstrated a rapid rise in neutralizing titers between d5-15 that plateaued ∼1:250 and remained elevated through the time course tested (**Figure 1B**).

**Fig. 1.**
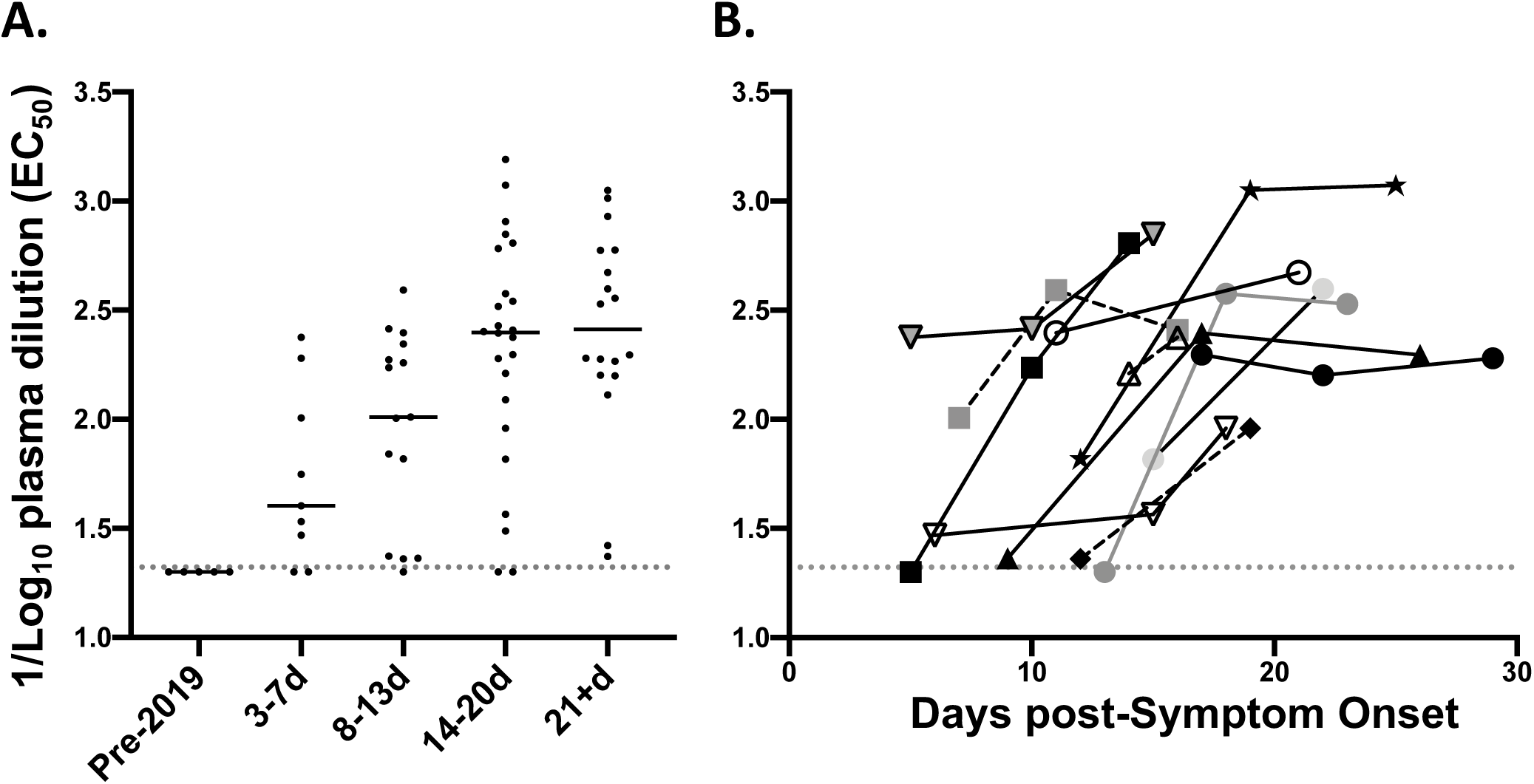
SARS-CoV-2 neutralizing titers in patients with and without PCR-confirmed COVID-19 Infection. **(A)** Neutralizing titers of 5 control specimens collected in 2015 and stored at -80°C and 67 specimens from 48 patients with PCR-positive COVID-19 relative to days from symptom onset. **(B)** Neutralizing titers relative to days of symptom onset. **(C)** Time to positive neutralizing antibodies in 12 patients with serial samples. Gray dotted horizontal lines represent the limit of detection at 1:20.

The correlation of the SARS-CoV-2 neutralizing titer with the ratio reported by the Roche, Abbott, and EI assays was 0.29, 0.47, and 0.46 respectively (**Figure 2A-C)**. Higher neutralizing titers were generally associated with a higher ratio as measured by all three assays. At a cutoff of 1:32 for the neutralizing assay, the concordance kappa with Roche was 0.61 (95% CI; 0.35-0.86), with Abbott was 0.65 (0.42-0.88), and with EI was 0.69 (0.49-0.89). For all three assays, the concordance decreased with an increased threshold for neutralizing titers.

**Fig. 2.**
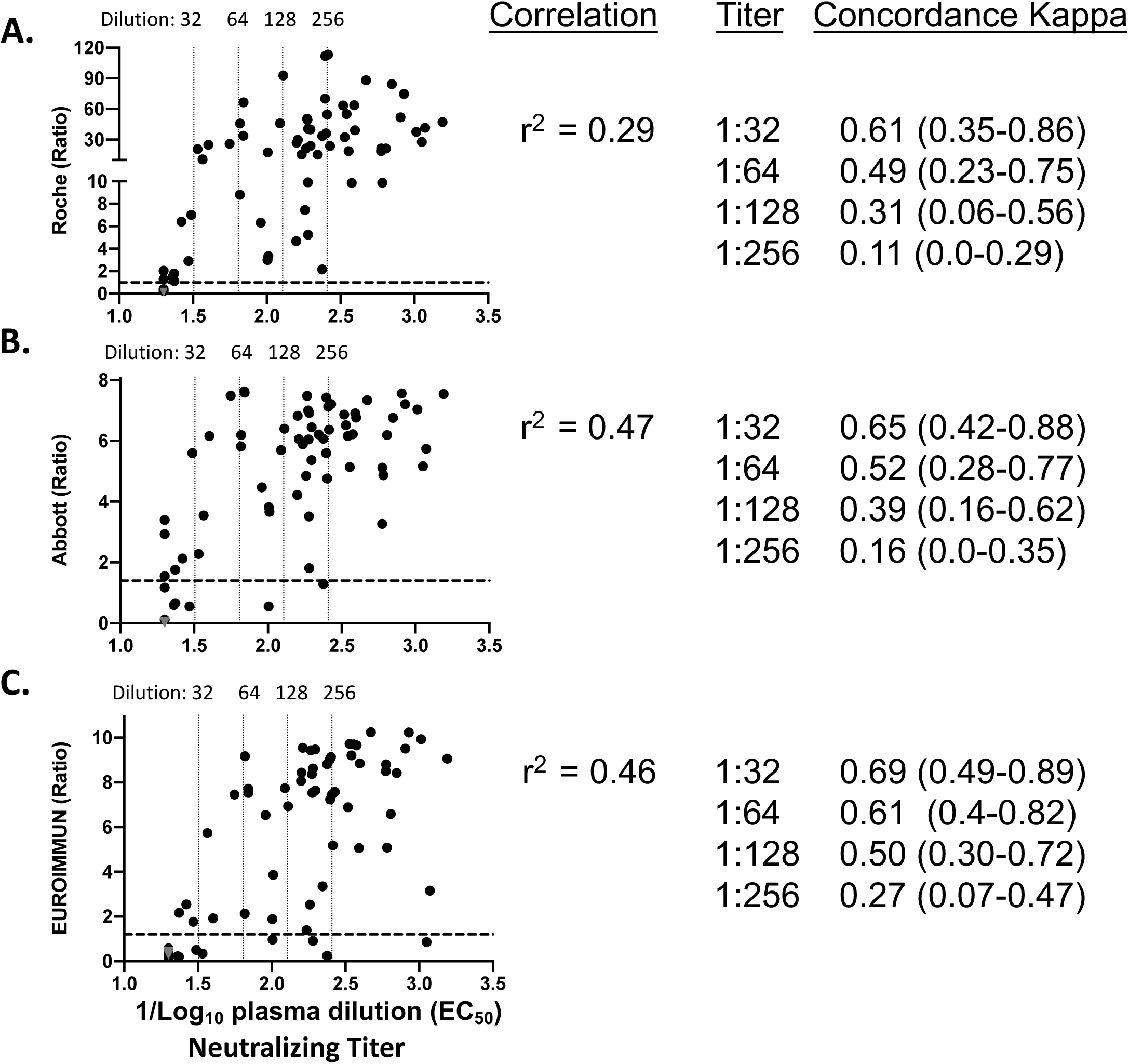
Correlation between neutralizing antibody titer and three commercial anti-SARS-CoV-2 serology assays. (A) Roche SARS-CoV-2 total antibody Immunoassay. Horizontal dotted line represents the cutoff off for Roche positivity (Ratio 1.0). (B) Abbott SARS-CoV-2 IgG Immunoassay. Horizontal dotted line represents the cutoff off for Abbott positivity (Ratio 1.4). (C) EUROIMMUN anti-SARS-CoV-2 IgG ELISA. Horizontal dotted line represents the cutoff off for EUROIMMUN positivity (Ratio 1.2). Specimens from 5 expected negative specimens collected in 2015 (gray triangles) and 67 specimens from 48 patients with PCR-positive COVID-19. Vertical dotted lines represented the cutoff for neutralizing antibody positivity at the indicated titer.

ROC curves to determine the PPA and NPA of a positive antibody result on commercial assays for neutralizing titers ≥ 1:32 revealed an AUC of 0.94 (95% CI; 0.88-1.0), 0.89 (0.79-0.99), and 0.93 (0.87-0.99) for the Roche, Abbott and EI assays respectively (**Figure 3A**). For both the Roche and Abbott assays, the ratio established by the manufacturers produced maximum PPA with decreased NPA for neutralizing antibodies. Lowering the cutoff for EI increased the PPA without negatively impacting NPA. When evaluated for a neutralizing titer of 1:128, the AUC of the Roche assay was 0.86 (95% CI;0.77-0.95), for the Abbott was 0.82 (0.71-0.94), and for the EI was 0.9 (0.83-0.97) (**Figure 3B**). At this neutralizing titer, the manufacturers’ ratios for a positive result for all three assays maximized PPA while reducing NPA for anti-SARS-CoV-2 neutralizing antibodies.

**Fig. 3.**
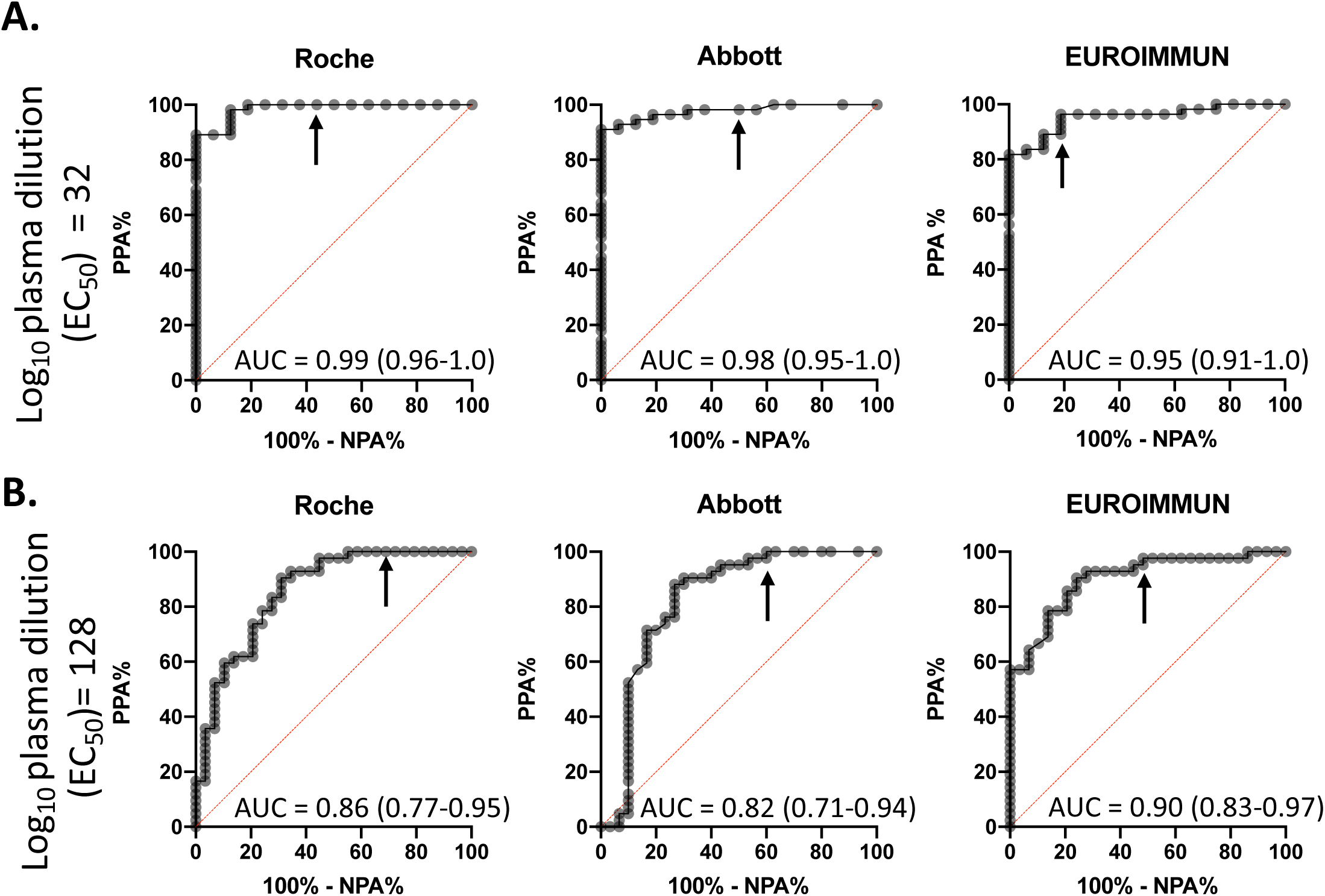
Receiver operating characteristic (ROC) curves for three commercial anti-SARS-CoV-2 serology assays to detect neutralizing anti-SARS CoV-2 antibodies. (A) Titer for neutralizing antibody positivity set at EC_50_=32. (B) Titer for neutralizing antibody positivity set at EC_50_=128. Dotted line represents AUC 0.5 (random guess line). Specimens from 5 expected negative specimens collected in 2015 and 67 specimens from 48 patients with PCR-positive COVID-19. Arrows represents commercial assay cutoff (Roche Ratio= 1.0; Abbott Ratio = 1.4; EUROIMMUN Ratio = 1.2). AUC= area under the curve.

At a neutralizing titer of 1:32, the PPA and NPA for the Roche assay was 100% (95% CI; 94-100) and 56% (30-80) at a ratio of 1.0 (**Table 1**). The ratio for each assay that improved the NPA while minimally affecting the PPA was assessed. The NPA improved to 81% (54-96) with the same PPA if the ratio for a positive result on the Roche was increased to 2.1. For the Abbott assay, the PPA was 96% (88-99) and the NPA was 69% (44-86) at a ratio of 1.4. The PPA and NPA for the Abbott changed to 95% (85-99) and 88% (65-96) respectively if the ratio for a positive result was adjusted to 2.2. For the EI assay, the PPA was 91% (80-96) and the NPA was 81% (57-93) at a cutoff of 1.2. By decreasing the ratio for a positive result to 0.72, the PPA improved to 96% (88-99) without effecting the NPA. NPA decreased for all three assays with increasing cutoff for a protective titer. To achieve an NPA >70% for all three assays at a neutralizing titer of 1:128, the ratio for a positive result would be 13.0 for the Roche, 4.8 for the Abbott, and 2.4 for the EI assays. PPA remained above 80% for all assays at these cutoffs.

**TABLE 1.** PPA and NPA of SARS-CoV-2 serological assays for neutralizing antibodies at multiple neutralizing titers

| Neutralizing<br>Titer |  | Roche |  |  | Abbott |  |  | EUROIMMUN |  |  |
| --- | --- | --- | --- | --- | --- | --- | --- | --- | --- | --- |
|  |  | Ratio | PPA<br>(95% CI) | NPA<br>(95% CI) | Ratio | PPA<br>(95% CI) | NPA<br>(95% CI) | Ratio | PPA<br>(95% CI) | NPA<br>(95% CI) |
| 1:20 | Manufacturer<br>Ratio | 1 | 100 (94-100) | 69 (42-87) | 1.4 | 93 (84-97) | 69 (42-87) | 1.2 | 90 (79-85) | 92 (67-100) |
|  | Ideal Ratio | 2.1 | 100 (94-100) | 100 (77-100) | 3.5 | 86 (73-92) | 100 (77-100) | 0.72 | 95 (86-99) | 92 (67-100) |
| 1:32 | Manufacturer<br>Ratio | 1.0 | 100 (94-100) | 56 (30-80) | 1.4 | 96 (88-99) | 69 (44-86) | 1.2 | 91 (80-96) | 81 (57-93) |
|  | Ideal Ratio | 2.1 | 100 (94-100) | 81 (54-96) | 2.2 | 95 (85-99) | 88 (65-96) | 0.72 | 96 (88-99) | 81 (57-93) |
| 1:64 | Manufacturer<br>Ratio | 1.0 | 100 (93-100) | 47 (27-68) | 1.4 | 96 (87-99) | 50 (30-70) | 1.2 | 92 (82-97) | 70 (48-85) |
|  | Ideal Ratio | 3.0 | 98 (90-100) | 74 (51-88) | 2.6 | 94 (84-98) | 70 (48-85) | 0.72 | 98 (90-100) | 70 (48-85) |
| 1:128 | Manufacturer<br>Ratio | 1.0 | 100 (92-100) | 31 (17-49) | 1.4 | 98 (85-99) | 40 (25-58) | 1.2 | 95 (84-99) | 55 (38-72) |
|  | Ideal Ratio | 13.0 | 83 (69-92) | 72 (54-85) | 4.8 | 86 (72-93) | 73 (56-86) | 2.4 | 93 (81-98) | 72 (54-85) |
| 1:256 | Manufacturer<br>Ratio | 1.0 | 100 (85-100) | 18 (10-31) | 1.4 | 100 (85-100) | 24 (15-38) | 1.2 | 100 (85-100) | 35 (23-49) |
|  | Ideal Ratio | 28.0 | 68 (47-84) | 71 (58-82) | 6.1 | 77 (57-90) | 73 (60-84) | 7.6 | 68 (47-84) | 71 (58-82) |

Patients that died as a result of COVID-19 had higher neutralizing antibody titers (mean, 1:576) compared to patients that survived (mean, 1:162) (**Figure 4A**). In contrast, no significant difference in ratio was observed between patients that died from COVID-19 compared to those that survived using the Roche, Abbott, or EI assays. Increased neutralizing antibody titers were also higher in patients that were intubated, had cardiac injury, or AKI relative to those with milder COVID-19 symptoms (**Figure 4B-D**). In contrast, no significant differences were noted between the groups regardless of outcomes when using the Roche, Abbott, and EI assays. However, similar non-significant trends (*i*.*e*., increase in ratio) were observed in patients who were intubated, had cardiac injury, or AKI with the EI assay. Neutralizing titers trended higher in male patients and patients >60 years old, although this was not statistically significant. Similar trends were observed with the serology assay ratios as well (**Supplemental Figure 3**). If categorized by low (<1:256) or high neutralizing titers (>1:256), there were no significant differences in outcomes between patients. However, there was an increase in the ratio observed in high neutralizing titer patients (6.3, 95% CI; 5.7-6.9) compared to low titer patients (5.1, 95% CI; 4.1-6.1) on the Abbott assay and the EI assay (8.2, 7.1-9.2 vs. 6.1, 4.6-7.6) (**Supplemental Table 1**). A similar, but non-significant trend was observed with the Roche assay.

**Fig. 4.**
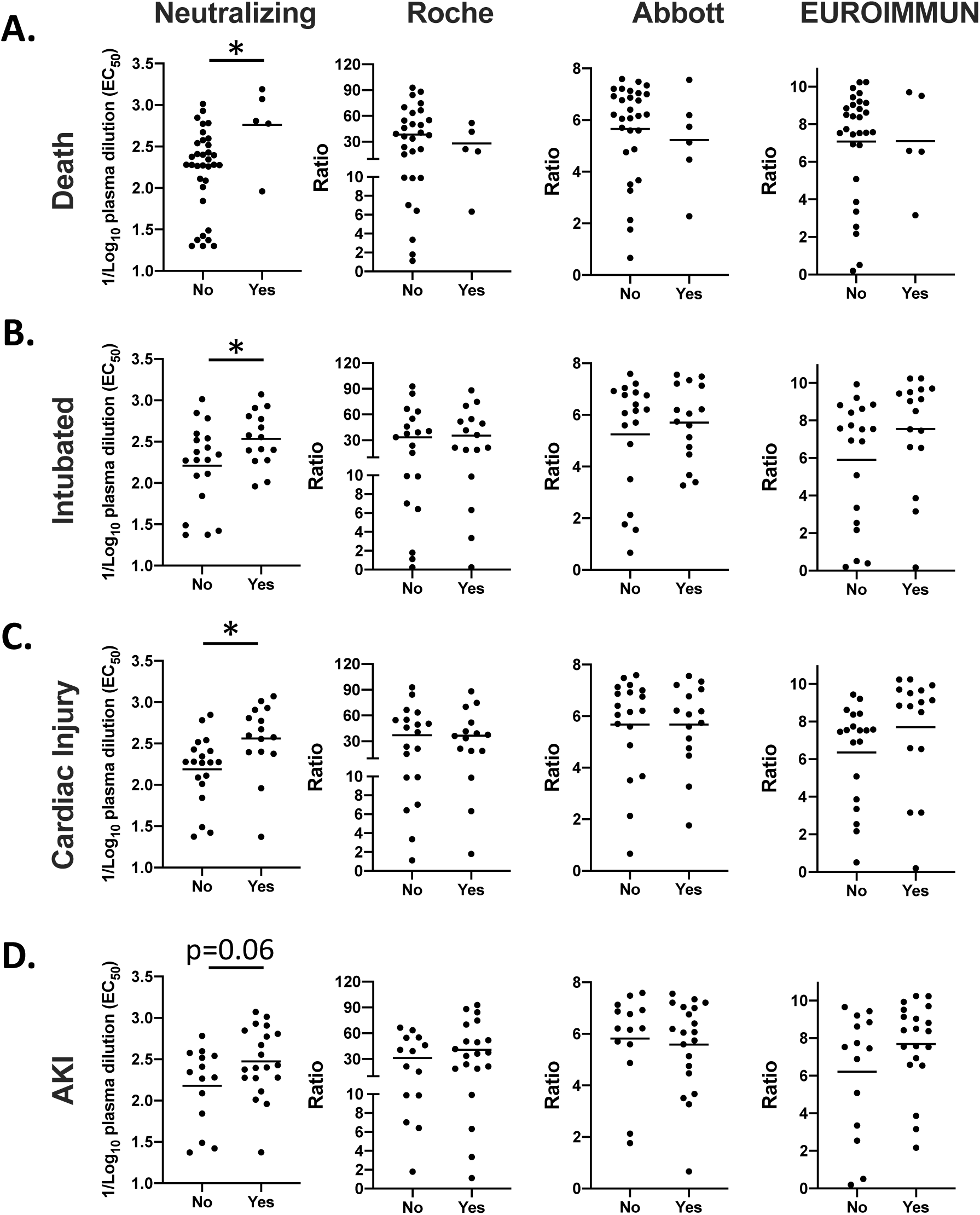
Association between clinical outcomes and anti-SARS CoV-2 neutralizing or commercial antibodies. (A) Death. (B) Intubation. (C) Cardiac Injury. (D) Acute kidney injury. Data from 40 patients with PCR-positive COVID-19. Solid horizontal line represents the mean. * p <0.05.

## DISCUSSION

The emergence of commercially available serological assays for the detection of antibodies to SARS-CoV-2 has outpaced scientific understanding of their immunological meaning and their value in clinical decision making. Here, we assessed the utility of three commercially available clinical assays for correlation with neutralizing antibodies to SARS-CoV-2. We observed modest correlation, but poor concordance and NPA between the Roche, Abbott and EI SARS-CoV-2 assays for the detection of SARS-CoV-2 neutralizing antibodies. Interestingly, the three commercial assays demonstrated similar performance with modest correlation but poor concordance and NPA for the detection of neutralizing antibody titers. Several studies have demonstrated that neutralizing antibodies are primarily against the S1, S2, and RBD domains of the SARS-CoV-2 spike protein *(3, 4)*. As a result, clinical assays targeting these regions have been hypothesized to better predict neutralizing titers. However, our findings indicate that the Roche (nucleocapsid), Abbott (nucleocapsid), and EI (S1) assays have similar performance for identifying patients with neutralizing antibodies. This implies that patients infected with SARS-CoV-2 develop a broad-based antibody repertoire against multiple proteins and epitopes, with a relatively fixed proportion of those acting as neutralizing antibodies.

While the World Health Organization (WHO) and the Centers for Disease Control (CDC) have advised against associating immunity with seropositivity *(20, 21)*, some have proposed that this warning is unnecessarily conservative *(22)*. Our findings suggest that SARS-CoV-2 serological assays should be interpreted with caution. While the majority of patients with antibodies detected by commercial assays had neutralizing antibodies present by d14 post-onset of symptoms, ∼10% of patients past d14 had titers that were <1:32. This implies that some patients with previous SARS-CoV-2 infections and positive antibody results by commercial assays may have neutralizing antibodies near the cutoff for a positive result. Although further studies are warranted, these low titers may be inadequate for protection, particularly if neutralizing antibodies are the primary therapeutic benefit of convalescent plasma. While higher reported ratios from all three commercial assays correlated with higher neutralizing titers, this was not universally true. Consistent with this, the correlation between neutralizing titers and serological results were <0.5 on all three commercial assays. These findings are consistent with a previous study demonstrating modest linear correlation between neutralizing SARS-CoV-2 titers with anti-RBD IgG or anti-S IgG using laboratory developed ELISAs *(23)*. Nonetheless, we found that higher ratios reported by all three commercial assays was associated with higher neutralizing titers. Importantly, all three serological assays used in this study currently have Emergency Use Authorization (EUA) to qualitatively determine the presence of antibodies against SARS-CoV-2. While a negative result on SARS-CoV-2 serological assays is likely to be associated with the absence of neutralizing antibody titers, a positive result is not reliable for predicting the presence of neutralizing antibodies. Furthermore, since these assays are under the EUA, they cannot be modified by the laboratory to report quantitative units. Our results argue for a potential utility in reporting the ratio calculated for commercially available assays relative to the calibrator. We, along with others, have previously suggested that commercially available serological assays for SARS-CoV-2 may have utility for identifying convalescent plasma donors *(24, 25)*. To this end, reporting quantitative units is more likely to identify convalescent patients with higher neutralizing antibody titers than qualitative cutoffs. Furthermore, if neutralizing antibodies are shown to confer protection to SARS-CoV-2, quantitative serological assays may assist in identifying neutralizing titers in mildly symptomatic and asymptomatic populations. However, further studies are needed to demonstrate the clinical benefit of this approach, especially by characterizing this association in a more diverse patient population.

While the NPA for neutralizing antibodies was >90% for all three commercial assays, this was only when a 1:20 neutralizing titer was used as a cutoff. It is important to note that this is far below the FDA recommended neutralizing titer for convalescent plasma donors (≥1:160) *(26)*. At a similar cutoff of 1:128, the NPA for neutralizing titers was below 60% for all three of the assays. Furthermore, while it is expected that neutralizing antibodies confer some protection against SARS-CoV-2, the titer required for this protective effect has not been established *(11)*. Due to the low sensitivity of serological assays for diagnosing early SARS-CoV-2 infection *(15, 27)*, some studies have suggested lowering the assay cutoff ratios to improve sensitivity *(28, 29)*. However, if the intended utility of serology is to determine the presence of neutralizing antibodies, our ROC analyses suggest that the assay cutoff should be increased to improve the NPA. Interestingly, some manufacturers are now associating positive serological results with neutralizing antibodies in their validation studies. For instance, the LIAISON SARS-CoV-2 S1/S2 IgG assay claims high agreement with neutralizing antibodies. However, the cutoff titer used for the neutralizing assay was 1:40; far below that recommended by FDA for convalescent plasma therapy *(13)*. If neutralizing antibodies >1:256 are required for protection, then commercial assays at the current cutoffs may have limited utility for identifying patients with protective antibodies; with NPA between 18-40% for the assays tested in this study.

Here, we observed that higher neutralizing titers are associated with worse clinical outcomes, a finding that was not observed with commercial serological assays. While seemingly counterintuitive, it is consistent with previous literature and may be a result of higher antigen burdens or hyperactive immune responses among other reasons *(30-34)*. A study of service members in the US Navy with predominantly mild symptoms revealed that ∼40% of those with a positive ELISA by the CDC assay had no neutralizing titers at a cutoff of 1:40 *(35)*. Similarly, a recent study demonstrated neutralizing titers at <1:50 in 33% of recovered patients and below 1:1000 in 79% of patients *(23)*. Our findings are also consistent with a study assessing the agreement between the EI IgG result and neutralizing titers on predominantly non-hospitalized convalescent plasma donors *(33)*. The authors demonstrated that at a neutralizing titer of 1:320, the PPA and NPA were 96% and 32% respectively and that neutralizing titers were higher in a small cohort of hospitalized patients. Similarly, we demonstrate higher neutralizing titers among patients with worse outcomes in an almost entirely hospitalized cohort. Unique to this study, we also compare commercial tests head-to-head and, by extension, compare serologies to two different protein antigens with similar results. Taken together, previous studies coupled with the findings presented here are consistent with the notion that neutralizing antibodies, while an important component of the immune response, *(3, 4)* are unlikely to be the only mechanism of SARS-CoV-2 clearance and protection. Other immune responses such as cellular immunity, T cells, antibody mediated cellular immunity and antibody mediated complement fixation likely play a pivotal role in protection from SARS-CoV-2.

Due to both heavy marketing and misunderstanding of their utility, patients have sought antibody testing for SARS-CoV-2 to determine if they had been previously infected and for peace-of-mind, assuming that they may have some level of protection (the concept of an “immunity passport”). At our institution, ∼85% of the SARS-CoV-2 serological tests are performed in the outpatient setting. This implies that the vast majority of these tests may be performed on mildly symptomatic and asymptomatic populations. Therefore, it is crucial that future studies address the correlation between neutralizing titers and commercial assays in the mildly symptomatic and the asymptomatic COVID-19 population. If symptomatic and severely ill patients have the highest titers of neutralizing antibodies, low concordance demonstrated here may be exacerbated by including asymptomatic and mildly symptomatic patients. Furthermore, while neutralizing titers appear to persist in the small group of patients with longitudinal specimens, the duration of follow up in our study was too short to determine the durability of neutralizing antibodies. Nonetheless, previous studies have demonstrated a reduction in neutralizing titers after 8 weeks post-hospital discharge *(31)*.

There are several limitations associated with this study. The true sensitivity and specificity of neutralizing titers in PCR-confirmed SARS-CoV-2 infected patients could not be accurately determined because specimens were pre-selected for serological positivity by commercially available immunoassays. This approach was chosen given the highly manual nature of testing for neutralizing antibodies and the primary goal of comparing neutralizing antibody titers to commercial assays. Furthermore, while the neutralizing assay utilized is robust and reproducible, it has not been validated for clinical use. In contrast to other studies, this assay uses an infectious strain of SARS-CoV-2 as opposed to pseudotyped rhabodoviruses or lentiviruses that heterologously express the SARS-CoV-2 spike protein. Furthermore, the relatively small number of patients tested means that potentially subtle differences in PPA, NPA, and concordance between the three assays could not be distinguished as a result of wide, overlapping confidence intervals. Finally, while others have demonstrated that neutralizing titers appear as early as d10 post-onset of symptoms, it is possible that assessing patients at later time points (*i*.*e*., d28) would reveal a higher concordance. While the majority of patients tested serially had neutralizing titers that peaked by d14-15, future studies are needed at later timepoints to correlation with commercial assays at later timepoints. This includes several months after infection, when other studies have demonstrated the neutralizing response beginning to diminish.

In conclusion, our findings suggest that positive serological results by three commercially available assays that measure antibodies against the viral spike or nucleocapsid protein of SARS-CoV-2 have modest correlation with neutralizing antibody titers. COVID-19 patients generate an antibody response to multiple viral proteins such that the quantitative ratios on the Roche, Abbott, and EUROIMMUN assays have comparable associations with SARS-CoV-2 neutralization. Nevertheless, commercial serological assays have poor NPA for SARS-CoV-2 neutralization, making them imperfect proxies for neutralization.

## Notes

### Competing Interest Statement

The authors have declared no competing interest.

