## Supplement for "Association between SARS-CoV-2 neutralizing antibodies and commercial serological assays"

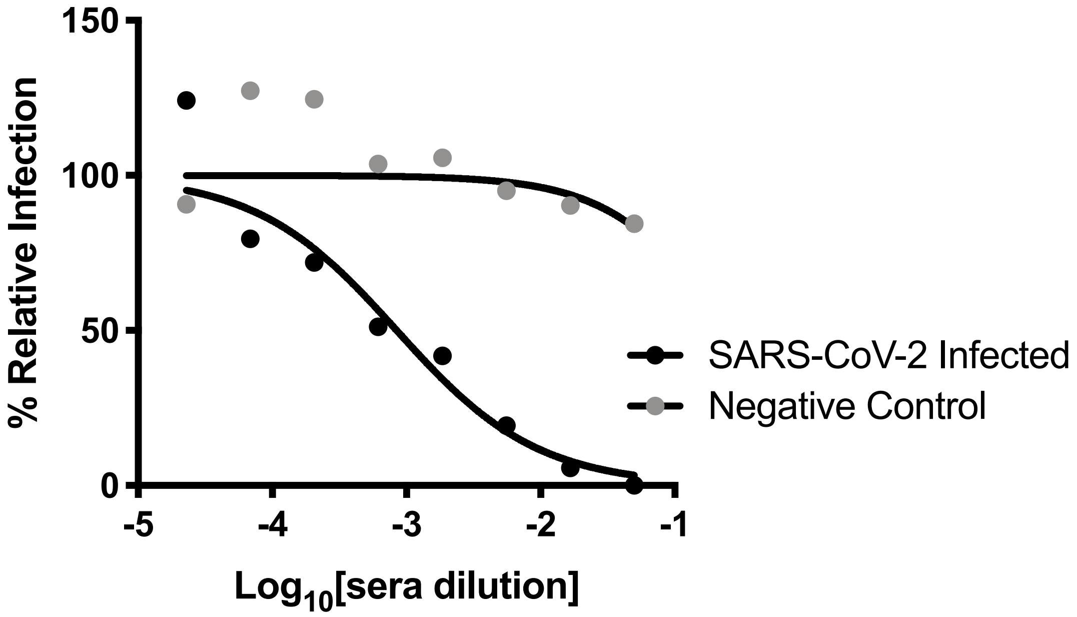


**Supplemental Figure 1. Serial dilution curves for SARS-CoV-2 neutralizing antibodies**. Black circles represent serum from a SARS-CoV-2 neutralizing antibody positive patient. Gray circles represent serum from a SARS-CoV-2 neutralizing antibody negative patient

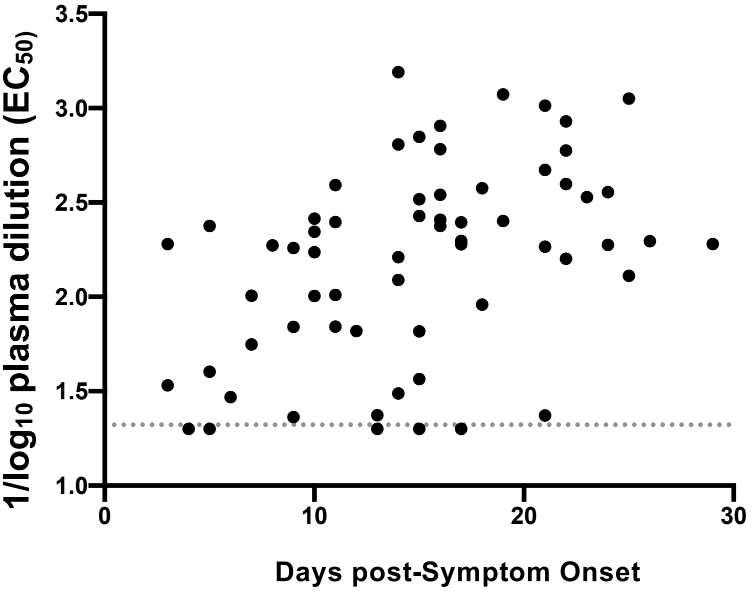


**Supplemental Figure 2. SARS-CoV-2 neutralizing titers in patients with PCR-confirmed COVID-19 Infection**. Neutralizing titers relative to days of symptom onset. Gray line is the limit of detection at 1:20.

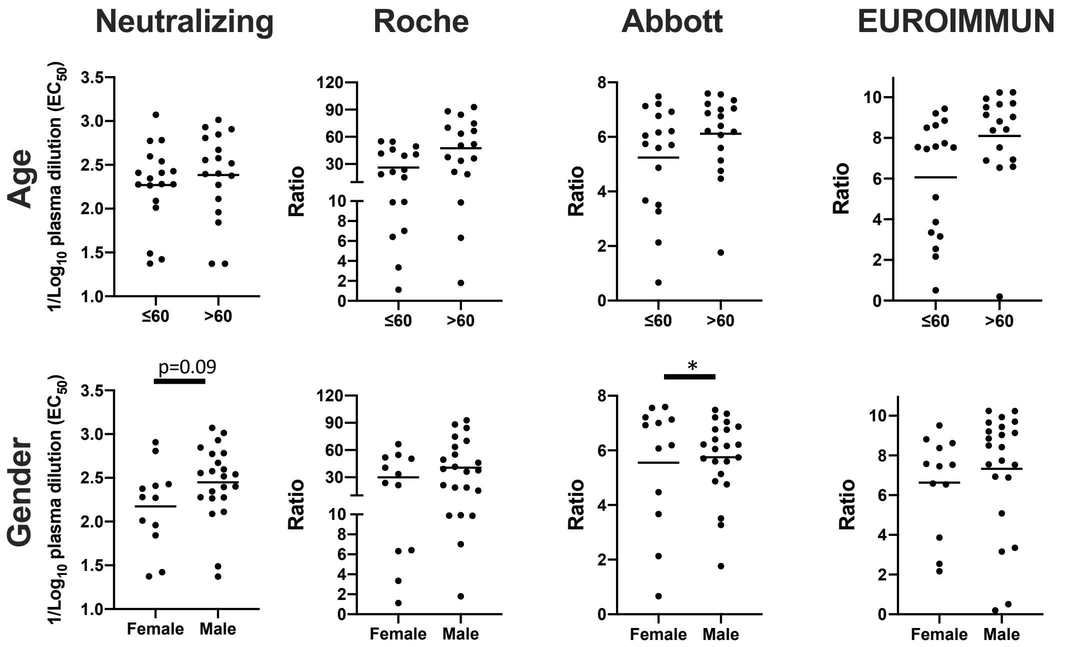


**Supplemental Figure 3. Association between age and gender with Anti-SARS CoV-2 neutralizing or commercial antibodies.** (A) Age ≤ 60 or > 60 and (B) gender. Solid horizontal line represents the mean. * $p < 0.05$.

**Supplemental Table 1: Association of high neutralizing titers with outcomes and commercial assay results**

| | **<1:256** | **>1:256** | ***p*** |
|---|---|---|---|
| n= | 18 | 16 | |
| Male Gender (%) | 10 (56) | 12 (75) | 0.3 |
| Age (95%) | 60 (52-67) | 58 (48-68) | 0.81 |
| Mortality (%) | 1 (5.6%) | 4 (25%) | 0.16 |
| Intubation (%) | 6 (33%) | 9 (64%) | 0.15 |
| Cardiac Injury (%) | 5 (29%) | 10 (63%) | 0.08 |
| AKI (%) | 10 (56%) | 10 (63%) | 0.73 |
| Roche* | 31.0 (17.4-44.6) | 43.3 (29.7-56.9) | 0.19 |
| Abbott* | 5.1 (4.1-6.1) | 6.3 (5.7-6.9) | **0.03** |
| EUROIMMIUN* | 6.1 (4.6-7.6) | 8.2 (7.1-9.2) | **0.03** |

*Results are mean (95% CI), p calculated from unpaired T test
All other p values calculated using Fischer's Exact test
